# Oligodendrocytes are a lifelong source of nuclear and ribosomal material for neurons in the mouse brain

**DOI:** 10.1101/2021.11.30.470658

**Authors:** Florian Mayrhofer, Angela M. Hanson, Carmen Falcone, Yang K. Xiang, Manuel F. Navedo, Wenbin Deng, Olga Chechneva

## Abstract

Nuclear and ribosomal components define cell identity and function by regulating chromatin dynamics, gene expression, and protein turnover. Here we report that in the mouse central nervous system (CNS) under normal conditions, neurons accumulate nuclear and ribosomal material of oligodendrocyte (OL) origin. We show that neuronal accumulation of OL-derived nuclear and ribosomal material is brain area-specific, and in the cortex and hippocampal dentate gyrus gradually propagates during postnatal brain maturation. We further demonstrate that OL-to-neuron material transfer persists throughout adulthood and responds to neuroinflammation. We found that satellite OL of the gray matter form internuclear contacts with receiving neurons in the mouse brain. Similar close internuclear associations between satellite OL and neurons are present in the adult human cortex. Our findings provide the first evidence of wide-spread dynamic and selective OL-to-neuron nuclear and ribosomal material transfer in the mouse CNS and indicate that satellite OL serve as powerful mediators of neuronal function. Equivalent processes may occur in the human CNS and cause neurological disorders when dysregulated.

**One Sentence Summary:** Neurons receive OL-derived nuclear and ribosomal material

## Main

Brain plasticity and cognitive function depend on the dynamic chromatin landscape and protein composition in neurons^1–5^. We used the Cre-LoxP system for the expression of individual ribosomal and nuclear reporter proteins, including the inner nuclear membrane protein Sun1 fused to superfolder green fluorescent protein (Sun1-sfGFP)^6^, the nuclear pore complex component RanGAP1 fused to mCherry (RanGAP1-mCherry)^7^, the core histone H2B fused to mCherry (H2B-mCherry)^7,8^, ribosomal subunit Rpl10a fused to enhanced green fluorescent protein (Rpl10a-EGFP)^9^ and ribosomal subunit Rpl22 fused to Hemagglutinin (Rpl22-HA)^10^ in the validated oligodendrocyte (OL) Cre mouse line, *SOX10-Cre*^11,12^. Unexpectedly we found the presence of nuclear and ribosomal reporter proteins not only in OL but also in neurons throughout the entire gray matter of the adult central nervous system (CNS) (Fig. 1a-e, and Extended Data Fig. 1). Brain area-specific differences in the numbers of reporter-positive neurons was observed between cortex, thalamus, and striatum (Fig. 1d). Notable, in our control *SOX10-Cre*:*EYFP* mice, in which unbound EYFP reporter is expressed in OL, only low numbers of weakly fluorescent neurons (6.8% ± 1.5% in the cortex, 1.7% ± 0.9% in the thalamus, and 2.0% ± 1.0% in the striatum) were found (Extended data Fig. 2a-e), as described earlier^13^. We did not see the presence of reporter proteins in microglia, a dominant phagocytic cell type in the CNS^14^ (Extended Data Fig. 3). Some reporter-positive astrocytes were detected in the hippocampus and deep gray matter (DGM) including striatum, thalamus and hypothalamus (Extended Data Fig. 3).

**Fig. 1.**
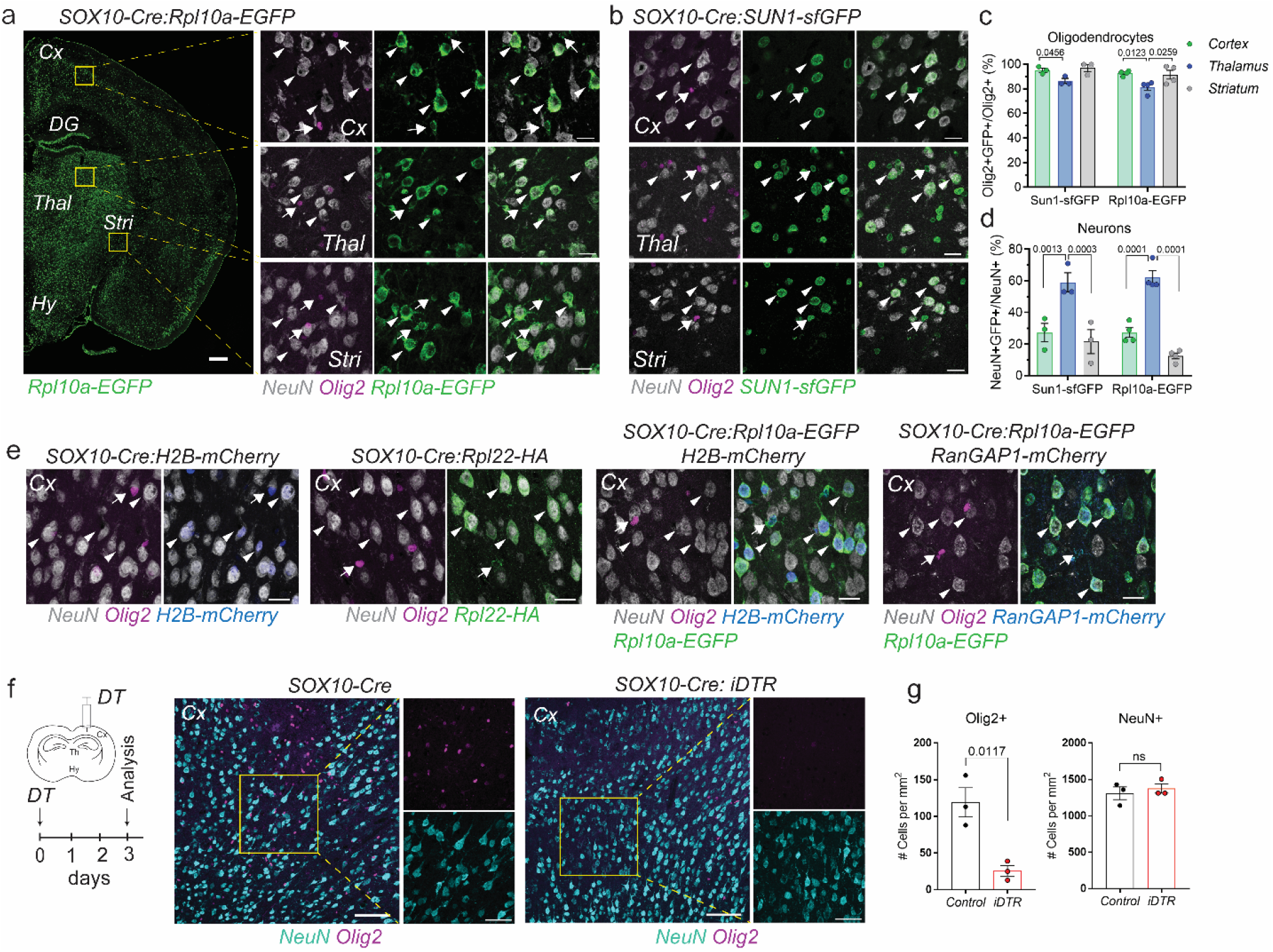
Neurons accumulate OL-derived nuclear and ribosomal material. (**a-b**) In the brain of adult *SOX10-Cre:Rpl10a-EGFP* and *SOX10-Cre:Sun1-sfGFP* mice, fluorescent reporters are localized to OL (arrow) and selective neurons (arrowheads). (**c-d**) Quantification of Olig2+GFP+ OL and NeuN+GFP+ neurons in the cortex, thalamus and striatum of adult *SOX10-Cre:Rpl10a-EGFP* and *SOX10-Cre:Sun1-sfGFP* mice. Data are presented as mean ± sem. Each circle represents an individual mouse. *P* values were determined by One-way ANOVA with Bonferroni post hoc test. (**e**) OL-derived nuclear and ribosomal material in OL (arrow) and selective neurons (arrowheads) in the cortex of adult *SOX10-Cre:H2B-mCherry, SOX10-Cre:Rpl22-HA, SOX10-Cre:Rpl10a-EGFP H2B-mCherry, SOX10-Cre:Rpl10a-EGFP RanGAP1-mCherry* mice. (**f**) Ablation of OL in the cortex of *SOX10-Cre:iDTR* mice 3 days after DT injection. *SOX10-Cre:iDTR* and control *SOX10-Cre* mice received stereotaxic injection of 1 ng DT into the cortex. Neuronal density at the injection site in *SOX10-Cre:iDTR* mice is indistinguishable from neuronal density in control *SOX10-Cre* mice. (**g**) Quantification of Olig2+ OL and NeuN+ neurons in the cortex of *SOX10-Cre* and *SOX10-Cre:iDTR* mice. Data are presented as mean ± sem. Each circle represents an individual mouse. *P* values were determined by unpaired t-test. Scale bars 500 μm for a; 100 μm for f; 50 μm for enlargement in f; 20 μm for enlargements in a and for b, e. Cx, cortex; DG, dentate gyrus; DT, diphtheria toxin; DTR, diphtheria toxin receptor; HA, hemagglutinin; Hy, Hypothalamus; ns, not significant; Stri, striatum; Thal, thalamus.

To test for unexpected transient expression of Cre in neurons, we crossed *SOX10-Cre* mice with inducible diphtheria toxin receptor (iDTR) reporter mice. In the resulting offspring, Cre-dependent expression of iDTR mediates cell lineage ablation after diphtheria toxin (DT) administration^15^. Stereotaxic injection of DT to the cerebral cortex or thalamus of *SOX10-Cre:iDTR* mice resulted in the ablation of OL at the injection site 3 days after treatment (Fig. 1f-g, Extended Data Fig. 4). The presence of neurons with location and numbers indistinguishable from neurons in control *SOX10-Cre* mice at the injection site, supports our conclusion that in our transgenic system, neurons do not express reporter protein.

Most neurons are generated during embryonic development^16,17^. During postnatal maturation the brain undergoes rigorous remodeling of cortical and subcortical structures, dendritic growth, establishment of synaptic connections, and myelination^18–21^. We used *SOX10-Cre:Rpl10a-EGFP* mice to follow neuronal accumulation of OL-derived material in the postnatal brain. At postnatal day (P) 5, in the cortex or hippocampal DG very few neurons were positive for OL-derived reporter protein (Fig. 2), while multiple fluorescent neurons were present in the septum and thalamus early after birth (Extended Data Fig. 5a-b). The number of fluorescent neurons in the cortex and DG gradually increased by P30 (Fig. 2). We also found a gradual increase in reporter-positive Olig2+ OL in the cortex and DG (Extended Data Fig. 5c-d). We conclude that during postnatal brain maturation OL-to-neuron material transfer emerges in a regulated spatiotemporal pattern.

**Fig. 2.**
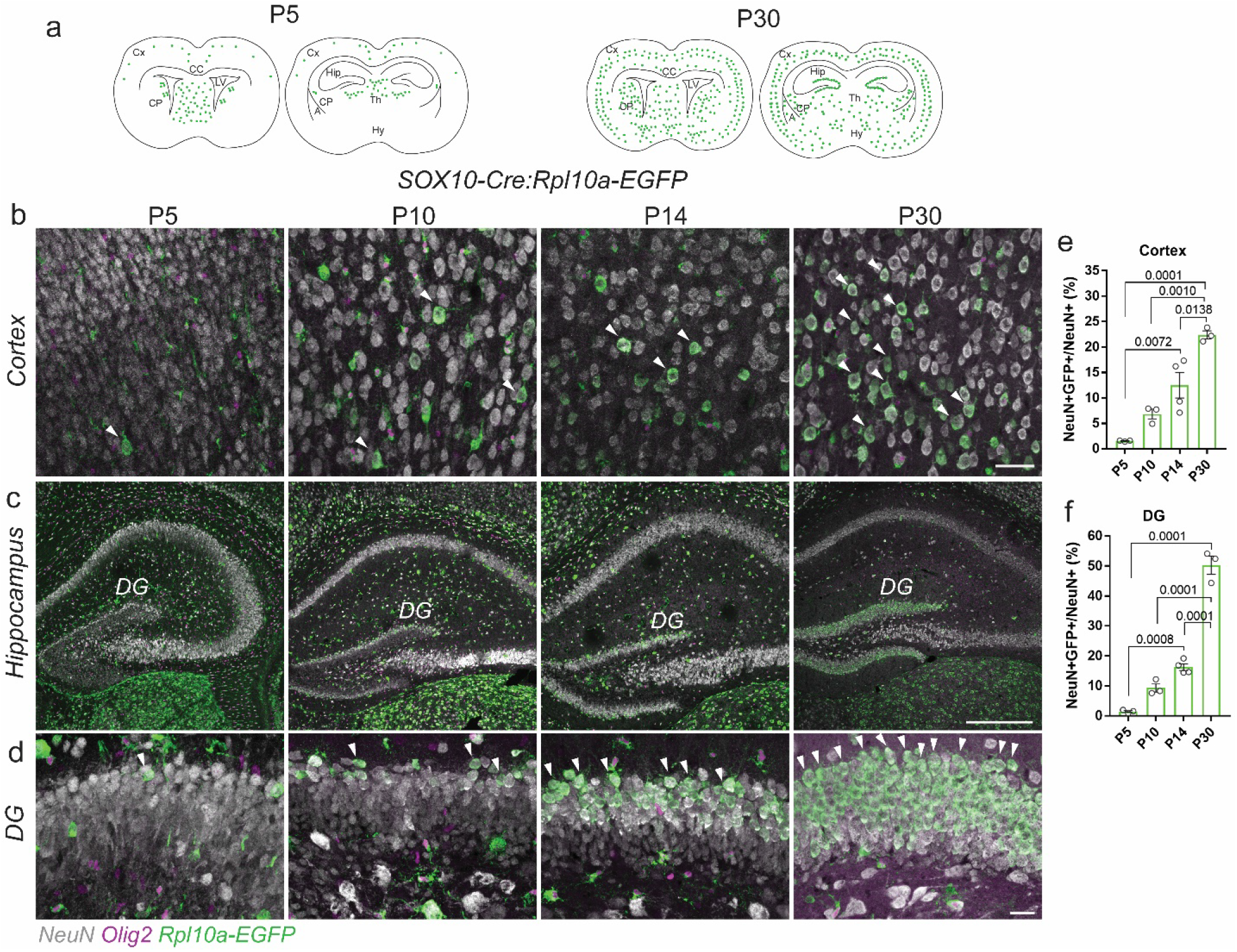
Postnatal propagation of OL-derived material in neurons. (**a**) Schematic showing the propagation pattern of fluorescent neurons in the cortex and hippocampal DG during postnatal development. (**b**) The number of Rpl10a-EGFP+ neurons (arrowheads) gradually increases from P5 to P30 in the cortex of *SOX10-Cre:Rpl10a-EGFP* mice. (**c-d**) Postnatal propagation of Rpl10a-EGFP+ neurons (arrowheads) in the hippocampal DG. (**e-f**) Quantification of Rpl10a-EGFP+ neurons in the cortex and DG. Data are presented as mean ± sem. Each circle represents an individual mouse. *P* values were determined by One-way ANOVA with Bonferroni post hoc test. Scale bar 500 μm for c; 50 μm for b; 20 μm for d. A, amygdala; CP, caudate putamen; CC, corpus callosum; Cx, cortex; DG, dentate gyrus; Hip, hippocampus; Hy, hypothalamus; P, postnatal day; Th, thalamus.

To study whether neurons receive nuclear and ribosomal material in the adult CNS, we employed the inducible CreERT2/LoxP system. In this system activation of CreERT2 recombinase by synthetic estrogen receptor ligand tamoxifen (TAM) allows controlled expression of reporter protein. We used the validated OL-specific mouse line *SOX10-iCreERT2*^12,22^. Four days after first TAM injection to *SOX10-iCreERT2:Sun1-sfGFP* or *SOX10-iCreERT2:Rpl10a-EGFP* mice, in addition to OL immunoreactive for nuclear or ribosomal reporter protein, we found reporter-positive neurons in the cortical layers 2-3 and 6, DG, septum, and amygdala (Fig. 3a-c, Extended Data Fig. 6a-c). Single fluorescent neurons were detected in the hippocampal CA1-3 pyramidal layer at this time point (Extended Data Fig. 6d). We also observed brain area-specific neuronal accumulation of OL-derived nuclear and ribosomal material in inducible OL lineage Cre mouse lines, *PDGFRα-iCreER*^23^ and *PLP-iCreER*^24^ (Extended Data Fig. 7). In *SOX10-iCreERT2* mice, Cre protein was specifically expressed in OL and none of the fluorescent neurons were immunoreactive for Cre (Fig. 3d), supporting the idea of selective OL-to-neuron material transfer in the adult brain.

**Fig. 3.**
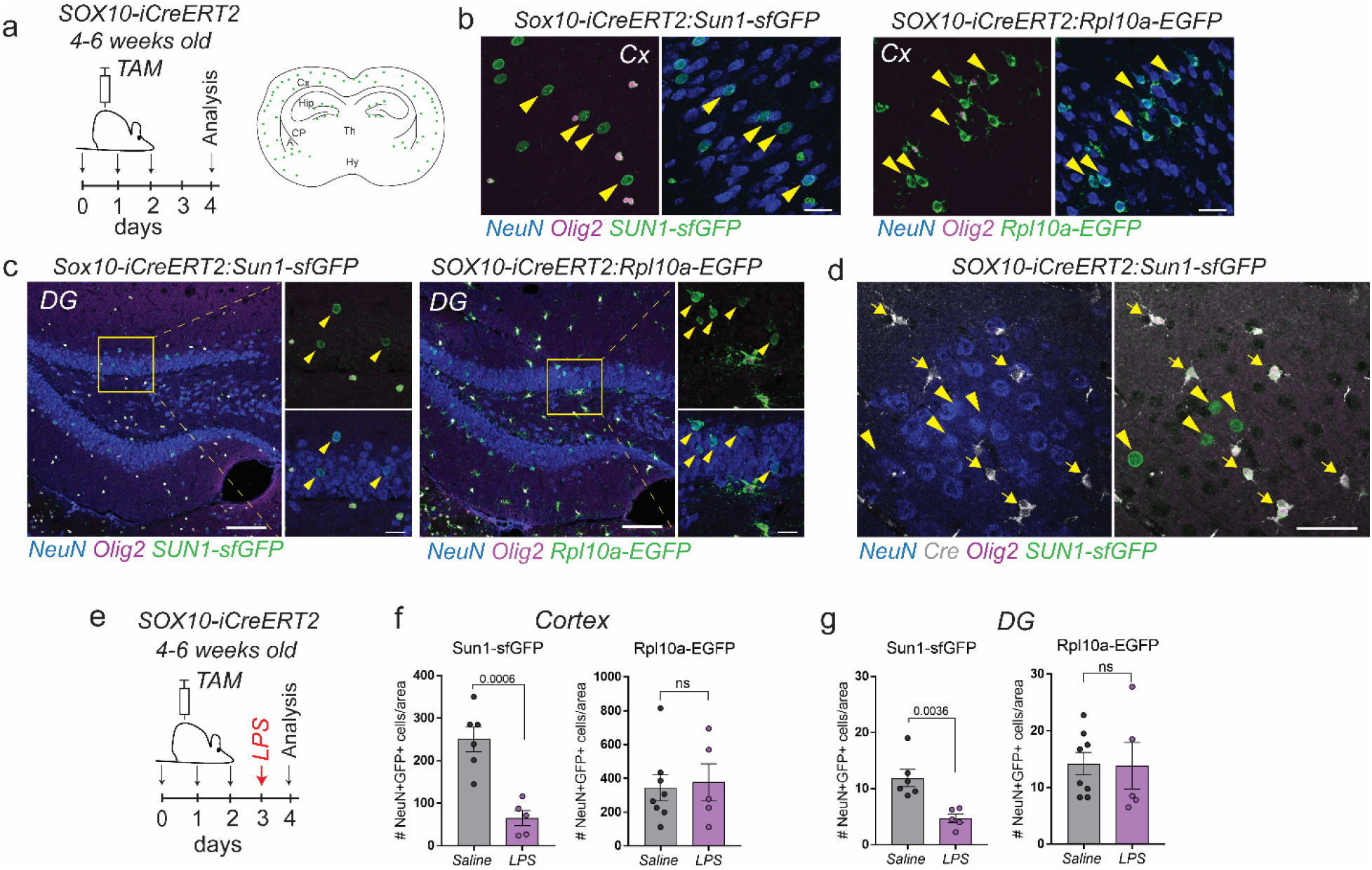
OL-to-neuron material transfer in the adult mouse brain. (**a**) Tamoxifen (TAM) was administered to 4-6 weeks old *SOX10-iCreERT2:Sun1-sfGFP and SOX10-iCreERT2:Rpl10a-EGFP* mice. At 4 days after first TAM injection fluorescent neurons were found in the cortical layers 2-3 and 6, hippocampal DG, and amygdala. (**b-c**) Fluorescent neurons (arrowheads) in the cortex and hippocampal DG of *SOX10-iCreERT2:Sun1-sfGFP and SOX10-iCreERT2:Rpl10a-EGFP* mice at 4 days post TAM. (**d**) Cre expression is restricted to OL (arrows) in the cortex of *SOX10-iCreERT2:Sun1-sfGFP* mice. No Cre expression is detected in fluorescent neurons (arrowheads). (**e**) LPS (4 mg/kg) was injected intraperitoneally 3 days post TAM to induce systemic inflammation. (**f-g**) Quantification of NeuN+GFP+ fluorescent neurons in cortical layers 2-3 and hippocampal DG in saline-and LPS-injected mice at 24h after injection. Data are presented as mean ± sem. Each circle represents an individual mouse. *P* values were determined by unpaired t-test. Scale bar 25 μm for b; 100 μm for c and 20 μm for enlargement in c; 50 μm for d. A, amygdala; CP, caudate putamen; Cx, cortex; DG, dentate gyrus; Hip, hippocampus; Hy, hypothalamus; LPS, lipopolysaccharide; ns, not significant; TAM, tamoxifen; Th, thalamus.

Neuroinflammation is a critical component of many neurological conditions including Alzheimer’s disease, cancer, multiple sclerosis, stroke, brain and spinal cord injury^25–27^. Neuroinflammation impairs glia-neuron interaction^27^. We found that acute neuroinflammation induced by endotoxin lipopolysaccharide (LPS) reduced the numbers of cortical and hippocampal DG neurons immunoreactive for nuclear reporter in *SOX10-iCreERT2:Sun1-sfGFP* mice 24 hours after LPS injection (Fig. 3e-g). No changes in accumulation of ribosomal reporter *Rpl10a-EGFP* were observed at this time point (Fig. 3f, g). We conclude that OL-to-neuron material transfer is a dynamic process that responds to injury.

In the gray matter, satellite OL (SOL) are uniquely positioned in intimate contact with the neuronal soma^28,29^. By examination of fluorescent neurons in the cortex of *SOX10-iCreERT2:Sun1-sfGFP* reporter mice four days after TAM induction, we detected immunoreactive SOL in close proximity to immunoreactive neuronal nuclei (Fig. 4a-b). Internuclear contact between immunoreactive SOL and neuronal nucleus was evident in the cortex of *SOX10-Cre:Rpl10a-EGFP H2B-mCherry* mice, where we found shared reporter H2B-mCherry across both nuclei (Fig. 4c). Super resolution confocal imaging revealed nuclear membrane contacts of SOL and neuronal nucleus and loss of nuclear membrane integrity at the contact site in the cortex of *SOX10-iCreERT2:Sun1-sfGFP* mice, suggesting internuclear exchange between SOL and selective neurons (Fig. 4d-e, Video 1). Notable, Cre recombinase and Olig2 immunoreactivity was confined to SOL nucleus and not detected in neurons (Fig. 4f). In many SOL-neuron internuclear associations, dense chromatin of SOL and dispersed chromatin of neuronal nucleus were evident (Fig. 4b-c). Sometimes we found close internuclear associations between two immunoreactive neuronal nuclei (Extended Data Fig. 8). Using fluorescence-activated nuclear sorting to quantify NeuN+Olig2+ double nuclei in the adult mouse brain, we identified differences in the numbers of SOL coupled to neuronal nuclei between cortex and DGM (Fig. 4g-i, Extended Data Fig. 9), supporting the idea of brain area-specific internuclear interaction between SOL and neurons in the mouse brain. Using immunofluorescent labeling for DAPI, NeuN and Olig2, we found internuclear associations between SOL and neurons in the postmortem adult human cortex, reminiscent of our findings on SOL-neuron internuclear contacts in the mouse brain (Fig. 4j, k).

**Fig. 4.**
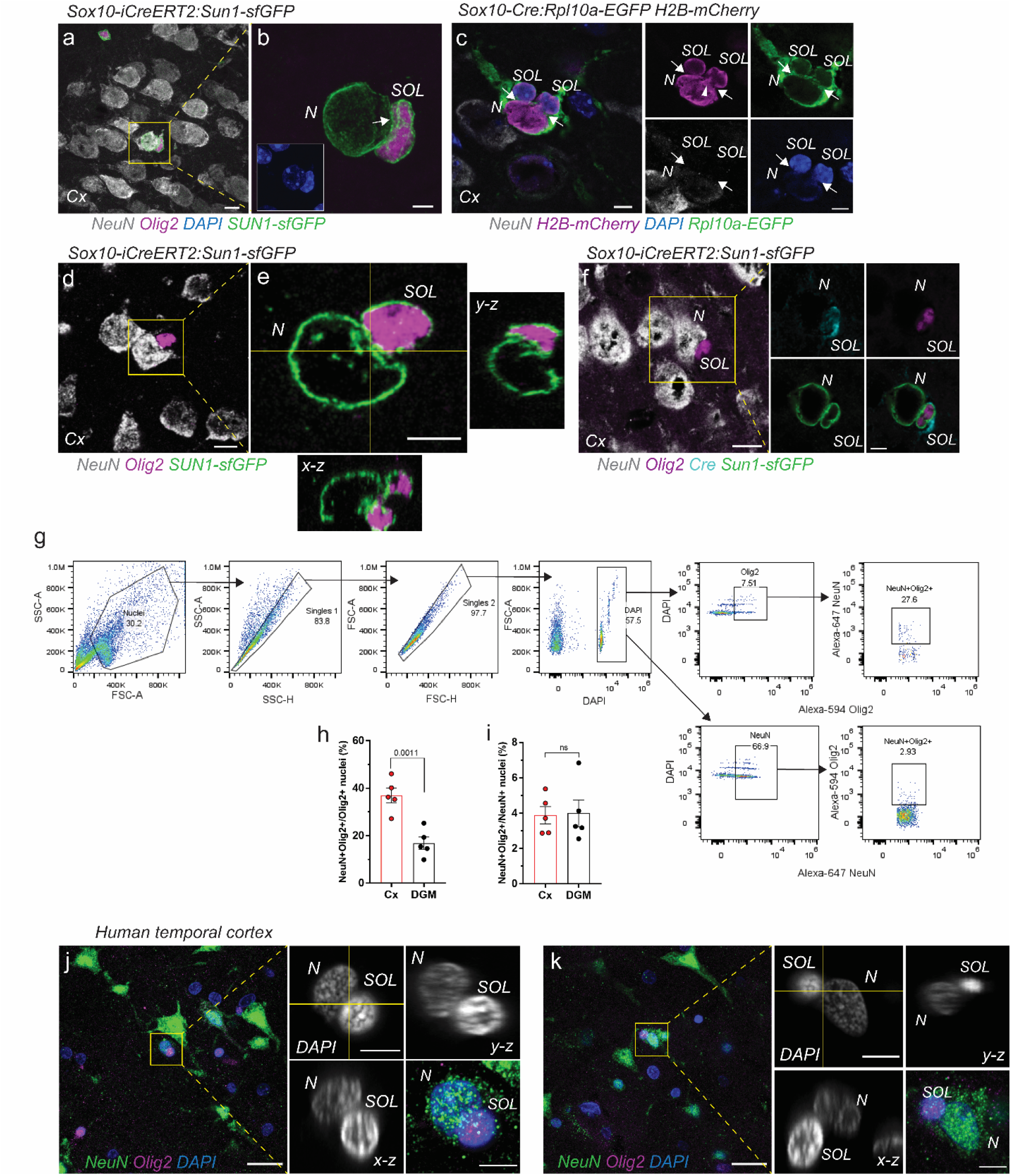
SOL and neurons form internuclear contact in the adult brain. (**a**) Confocal imaging of Sun1-sfGFP+ nuclear membrane reveals SOL in internuclear contact with selective neurons in the adult *SOX10-iCreERT2:Sun1-sfGFP* mouse cortex at 4 days post TAM. (**b**) Enlargement in A showing attachment of SOL nucleus to neuronal nucleus (arrow). Note DAPI-labeled dense chromatin of SOL and diffuse chromatin of attached neuronal nucleus. (**c**) Internuclear contacts between neuron and SOL (arrows) in the cortex of *SOX10-Cre:Rpl10a-EGFP H2B-mCherry* mouse. Note dense H2B-mCherry at internuclear contact site (arrowhead). (**d**) Super resolution confocal imaging of NeuN+ neuron and Olig2+ SOL internuclear couple in the cortex of adult *SOX10-iCreERT2:Sun1-sfGFP* mouse at 4 days post TAM. (**e**) Single plane super resolution image showing attachment of SOL and neuronal Sun1-sfGFP+ nuclear membranes and orthogonal projection at internuclear contact site. (**f**) Cre recombinase (cyan) and Olig2 (magenta) immunoreactivity were confined to SOL. (**g**) After isolation from the cortex and DGM of 12 weeks old *C57BL/6J* mice, NeuN+ and Olig2+ nuclei were analyzed using fluorescence-activated nuclei sorting. (**h**) Proportion of NeuN+Olig2+ nuclei in Olig2+ population. (**i**) Proportion of NeuN+Olig2+ nuclei in NeuN+ population. Data are presented as mean ± sem. Each circle represents an individual mouse. P value was determined by unpaired t-test. (**j-k**) Internuclear association between SOL and neuronal nucleus in the human temporal cortex of 68 years old male postmortem. Enlargements showing proximity of DAPI-labeled SOL and neuronal nucleus and orthogonal projection at attachment site. Note DAPI-labeled dense chromatin of SOL and diffuse chromatin of attached neuronal nucleus similar to the mouse brain. Scale bars 20 μm for j, k; 10 μm for a, d, f; 5 μm for b, c, e, and enlargement in f, j, k. Cx, cortex; DGM, deep gray matter; N, neuron; ns, not significant; SOL, satellite oligodendrocyte.

## Discussion

Here we demonstrate that in the mouse CNS under physiological conditions multiple neurons receive nuclear and ribosomal material from OL. While the notion of material transfer between OL and neurons has been previously postulated^30–33^, such widespread transfer of material as reported in our study is novel. Our data further suggest that nuclear and ribosomal material transfer is a selective and dynamic process that can be modulated.

OL comprise a highly heterogeneous glial cell population with thirteen distinct subtypes recently identified^34^. OL populations undergo dynamic brain region-specific changes during postnatal development^35^. OL heterogeneity, together with regional heterogeneity of neurons during postnatal maturation and in the adult brain^36^ may explain spatiotemporal differences of OL-to-neuron material transfer. Differences in the response to acute systemic inflammation between nuclear and ribosomal material transfer suggest differences in the underlying transfer mechanism or in material turnover inside neurons.

Myelinating OL support axonal transport and axonal metabolism by transcellular delivery of RNA and protein-carrying exosomes that are endocytosed at periaxonal sites^30,32,33^. The presence of internuclear contacts between SOL and neurons, reported here, suggests a more direct transfer of nuclear and ribosomal material from SOL to neurons. Confinement of Cre and OL-specific nuclear proteins such as Olig2 to OL and insignificant transfer of unbound EYFP indicates that material transfer is selective. Whether transferred material includes proteins, RNA or DNA and is packed into membrane-bound vesicles or organized in membrane-less nuclear condensates remains to be identified.

The biological significance of material transfer from OL to neurons reported here is largely unknown. However, given the importance of nuclear and ribosomal material in regulation of gene expression, protein synthesis, and cell function, our findings suggest a key role of OL in directly supporting neuronal activity. The presence of SOL-neuron internuclear pairs in the human brain suggests that a similar process of material transfer may also take place in the human CNS. The role of OL-to-neuron material transfer in CNS ribosomopathies^37^ or nuclear envelopathies^38^ or whether OL can be a source of pathological proteins that accumulate in neurons, such as amyloid-β and tau found in Alzheimer’s disease^39,40^, misfolded α-synuclein in Parkinson disease^41^, FUS and TAR DNA-binding protein 43 in frontotemporal dementia and amyotrophic lateral sclerosis^42,43^, and mutated huntingtin in Huntington disease ^44^ is unknown. Thus, elucidating the role of OL-to-neuron material transfer and targeting its mechanism to modulate neuronal function should prove to be an attractive field of exploration.

## Acknowledgments

We thank Dr. Laura Borodinsky, Dr. Nicholas Marsh-Armstrong, Dr. Fuzheng Guo, and Dr. Paul Knoepfler for critical reading of the manuscript and constructive suggestions. We thank Dr. Athena Soulika for constructive suggestions and technical support in nuclei sorting. We thank Jie Xu for technical support with the equipment. We thank Dr. Olga Balashova for critical reading of the manuscript, constructive suggestions, and technical advice in confocal imaging and image preparation. We thank Dr. David Pleasure for scientific support and access to the equipment.

## Author contributions

Finding, conceptualization and design, F.M., O.V.C.; Acquisition of data, F.M., O.V.C., A.M.H., M.F.N., C.F.; Resources and Equipment, W.D., Y.K.X.; Writing – Reviewing & Editing, O.V.C., F.M.

## Data and materials availability

All data is available in the main text and the supplementary materials.

---

**Video 1. SOL and neuron form internuclear contact in the adult mouse brain.** Animation through a Z-stack series of super resolution confocal imaging showing SOL-neuron internuclear contact in the cortex of *SOX10-iCreERT2:Sun1-sfGFP* mouse 4 days after TAM injection. Note Olig2+ SOL nucleus (magenta) and attached neuronal nucleus each enclosed by Sun1-sfGFP+ (green) nuclear membrane and loss of nuclear membrane integrity at the contact site.

## Methods

### Mice

Animals were maintained in accordance to the NIH Guide for the Care and Use of Laboratory Animals. Experimental protocols were approved by the Institutional Animal Care and Use Committee at the University of California, Davis.

Female or male reporter mice were bred with Cre positive mates. Offspring were genotyped for presence of the reporter transgene and Cre. Offspring carrying the reporter transgene and Cre positive were used as reporter animals in this study. Littermates carrying the reporter transgene and either Cre negative or non-induced CreERT2 positive were used as control. Adult mice used in the study were 4-12 weeks old. All transgenic lines used in this study are commercially available at The Jackson Laboratory.

#### SOX10-Cre

B6;CBA-Tg(Sox10-cre)1Wdr/J

Jax Stock No: 025807 (Donating Investigator: William Richardson, University College London). These mice express a nuclear-targeted Cre recombinase directed by the endogenous *SOX10* promoter/enhancer regions on a PAC transgene.

#### iSOX10-CreERT2

CBA; B6-Tg(Sox10-icre/ERT2)388Wdr/J

Jax Stock No: 027651 (Donating Investigator: William Richardson, University College London). These mice, carrying a transgene with tamoxifen-inducible optimized (improved) Cre recombinase under the control of the mouse Sox10 (SRY (sex determining region Y)-box 10) promoter, express CreERT2 in oligodendrocyte lineage cells.

#### EYFP

B6.129X1-Gt(ROSA)26Sortm1(EYFP)Cos/J

Jax Stock No: 006148 (Donating Investigator: Frank Costantini, Columbia University Medical Center). These mice have a LoxP-flanked STOP sequence followed by the Enhanced Yellow Fluorescent Protein gene (EYFP) inserted into the Gt(ROSA)26Sor locus. When bred to mice expressing Cre recombinase, the STOP sequence is deleted and EYFP expression is observed in the Cre-expressing tissues.

#### Rpl10a-EGFP (TRAP)

B6;129S4Gt(ROSA)26Sortm9(EGFP/Rpl10a)Amc/J

Jax Stock No: 024750 (Donating Investigator: Andrew P McMahon, University of Southern California). These mice have a transgene containing cDNA encoding enhanced green fluorescent protein (EGFP) fused to ribosomal protein unit L10a (Rpl10a) integrated into the Gt(ROSA)26Sor locus. Expression of the EGFP-tagged form of Rpl10a is blocked by a LoxP-flanked STOP fragment (3 copies of SV40 polyA) placed between Gt(ROSA)26Sor promoter and the Rpl10a-EGFP sequence. Cre-mediated excision of STOP fragment results in the constitutive expression of the Rpl10a-EGFP fusion gene.

#### Sun1-sfGFP

B6.129-Gt(ROSA)26Sortm5(CAG-Sun1/sfGFP)Nat/MmbeJ

Jax Stock No: 030952 or 021039 (Donating Investigators: Margarita Behrens, The Salk Institute; Jeremy Nathans, Johns Hopkins University). These Sun1-tagged mice are used in the INTACT (isolation of nuclei tagged in specific cell types) method for immunopurification of nuclei. Cre-dependent removal of a floxed STOP cassette allows expression of the SUN1 fusion protein at the inner nuclear membrane in targeted cell types.

#### iDTR

C57BL/6-Gt(ROSA)26Sortm1(HBEGF)Awai/J

Jax Stock No: 007900 (Donating Investigator: Ari Waisman, Johannes Gutenberg University of Mainz). These mice have the simian diphtheria toxin receptor (DTR; from simian Hbegf) inserted into the Gt(ROSA)26Sor (ROSA26) locus. Widespread expression of DTR is blocked by an upstream LoxP-flanked STOP sequence. The Cre-inducible expression of DTR in these mice render cells susceptible to ablation following Diphtheria toxin administration.

#### Rpl10a-EGFP RanGAP1-mCherry (NuTRAP)

B6; 129S6-Gt(ROSA)26Sortm2(CAG-NuTRAP)Evdr/J

Jax Stock No: 029899 (Donating Investigator: Evan D Rosen, Beth Israel Deaconess Medical Center and Harvard Medical School). These mice have a transgene containing cDNA encoding three individual components BirA, biotin ligase recognition peptide (BLRP)-tagged monomeric red fluorescent protein (mCherry) fused to mouse nuclear membrane Ran GTPase activating protein 1 (RanGAP1) and EGFP fused to ribosomal protein Rpl10a integrated into the Gt(ROSA)26Sor locus. Upon exposure to Cre recombinase, the NuTRAP allele co-expresses three individual components BirA, BLRP-tagged mCherry-RanGAP1 and Rpl10a-EGFP fusion gene.

#### Rpl10a-EGFP H2B-mCherry

B6.Cg-Gt(ROSA)26Sortm1(CAG-HIST1H2BJ/mCherry, EGFP/Rpl10a)Evdr/J

Jax Stock No: 029789 (Donating Investigator: Evan D Rosen, Beth Israel Deaconess Medical Center and Harvard Medical School). These mice have a transgene containing cDNA encoding human histone 1 H2bj gene (HIST1H2BJ) fused in-frame to the N-terminus of a monomeric mCherry and EGFP fused in-frame to the N-terminus of mouse 60S ribosomal subunit Rpl10a integrated into the Gt(ROSA)26Sor locus. Upon exposure to Cre recombinase, the H2B-TRAP allele co-expresses Rpl10a-EGFP and H2B-mCherry fusion gene.

#### H2B-mCherry

B6; 129S-Gt(ROSA)26Sortm1.1Ksvo/J

Jax Stock No: 023139 (Donating Investigator: Karel Svoboda, Janelia Farm Research Campus). These mice have a transgene containing cDNA encoding human histone 1 H2bb gene (HIST1H2BB) followed C-terminally by mCherry gene integrated into the Gt(ROSA)26Sor locus. Cre-mediated excision of the STOP fragment results in the constitutive expression of the H2B-mCherry fusion gene. The CNS tissue of H2B-mCherry Cre negative mice display mCherry background expression in all cells. As reported by the donating investigator, after exposure to Cre recombinase, the mCherry expression levels in the nucleus are significantly greater than those baseline levels.

#### RPL22-HA (RiboTag)

B6J.129(Cg)-Rpl22tm1.1Psam/SjJ

Jax Stock No: 029977 (Donating Investigator: Simon John, The Jackson Laboratory). These mice carry a targeted mutation of the ribosomal protein L22 (Rpl22) locus. Prior to Cre recombinase exposure, RiboTag mice express the wildtype RPL22 protein (15 kDa). Cre-mediated excision of the floxed exon 4 enables tissue-selective expression of the downstream HA epitope-tagged exon 4. The resultant 23 kDa HA epitope-tagged ribosomal protein (RPL22HA) is incorporated into actively translating polyribosomes.

#### PDGFRα-CreER

B6N.Cg-Tg(Pdgfra-cre/ERT)467Dbe/J

Jax Stock No: 018280 (Donating Investigator: Dwight E Bergles, Johns Hopkins School of Medicine). These mice express Cre-ER protein directed by the PDGFRα promoter/enhancer regions on a BAC transgene. Cre-ER can only gain access to the nuclear compartment after exposure to tamoxifen. However, in some PDGFRα-CreER:Rpl10a-EGFP mice we observed random aberrant Cre activity, where reporter transgene shows tamoxifen-independent expression in large numbers of neurons and astrocytes, as reported by donating investigator.

#### PLP-CreER

B6.Cg-Tg(Plp1-cre/ERT)3Pop/J

Jax Stock No: 005975 (Donating Investigator: Brian Popko, University of Chicago). These mice have a tamoxifen inducible Cre-mediated recombination system driven by the mouse proteolipid protein (PLP) 1 promoter on a transgenic construct.

### Tamoxifen induction of Cre activity

Tamoxifen (TAM, T5648 Sigma) was dissolved at the concentration of 20 mg/ml in corn oil by shaking at 37°C overnight protected from light. 4-6 weeks old mice received three intraperitoneal injections of 80 mg/kg TAM for three consecutive days. We observed Cre-induced recombination in non-injected Cre+ mice that were housed together with TAM injected littermates. Therefore, not-induced Cre+ control mice were housed separately from TAM injected cohorts.

### Stereotaxic intracerebral injection

Intracerebral injection of diphtheria toxin from Corynebacterium diphtheria (DT, D0564 Sigma) into the cerebral cortex or thalamus were performed using stereotaxic alignment system (David Kopf Instruments). In brief, 6-8 weeks old *SOX10-Cre:iDTR* and control *SOX10-Cre* mice were anesthetized with isoflurane and positioned in the stereotaxic frame. A 0.2 mm hole was drilled in the skull at coordinates of 1.0 mm posterior to bregma and 1.0 mm lateral to the midline in the right hemisphere. 1 ng of 1 ng/μl DT dissolved in artificial cerebrospinal fluid was injected at the depth of 1 mm below the surface of the skull for cortex and 3 mm for thalamus with a 33G needle attached to 5 μl Hamilton syringe. After 72 hours mice were transcardially perfused with ice cold phosphate buffered saline (PBS) followed by ice cold 4% paraformaldehyde (PFA), brains were isolated and processed for immunolabeling as described below.

### Neuroinflammation

To induce neuroinflammation, 4-6 weeks old mice received a single intraperitoneal (i.p.) injection of 4 mg/kg lipopolysaccharide (LPS) from E. coli O111:B4 (Sigma, L4391) or sterile saline 3 days after TAM induction. Mice were perfused 24 h after injection and brains were isolated and processed as described below.

### Immunolabeling and confocal imaging

Immunolabeling was performed on cryosections as previously described ^25^, with minimal modifications. Mice were anesthetized with isoflurane (5% for 6 min) and perfused with ice cold PBS followed by ice cold 4% PFA with flow rate 2 ml/min for 5 min each. Brains and spinal cords were dissected, postfixed in 4% PFA for 24 h and subsequently cryopreserved in 10% and 20% sucrose, embedded in OCT:20% sucrose 1:1 mixture in the vapor of liquid nitrogen. Tissues were cut in 15 μm sections on the cryostat. For immunolabeling, sections were washed 3 x 5 min with PBS to remove residues of OCT. Nonspecific binding of antibodies was blocked at room temperature (RT) for 30 min using 10% normal serum in 0.2% Triton-X in PBS. We used following antibodies: goat anti-mCherry (1:1000, Biorbyt orb11618), chicken anti-GFP (1:500, Novus Biologicals NB100-1614), rabbit anti-HA (1:50, Zymed 71-5500), goat anti-Olig2 (1:200, R&D AF2418), rabbit anti-Olig2 (1:200, Abcam AB109186), rabbit anti-NeuN (1:500, Millipore ABN78), mouse anti-NeuN (1:100, Millipore MAB377), mouse anti-Cre recombinase (1:200, Clone 2D2, Millipore MAB3120), rabbit anti-Iba1 (1:500; Wako 019-19741), rat anti-GFAP (1:500, Millipore 345860) diluted in PBS containing 5% normal serum and 0.2% Triton-X. Sections were incubated either for 2 h at RT or overnight at 4 °C. After 3 x 10 min wash with PBS, sections were incubated with fluorophore-conjugated secondary antibodies Alexa Fluor-405, 488, 594 or 647 (1:1000; Life Technologies) to detect primary antibody. Slides were mounted using DAPI Fluoromount-G (SouthernBiotech). For staining containing Alexa Fluor 405, slides were mounted using Fluoromount-G without DAPI.

Images were acquired on Nikon Eclipse 90i A1 and Nikon Eclipse Ti2 C2 laser scanning confocal microscopes. Negative controls omitting primary antibody were performed to confirm specificity of staining. In constitutive *SOX10-Cre* mice, cells were counted in low magnification images (x20 objective) of coronal sections in the cortex, thalamus, striatal caudate putamen, and hippocampal DG of 4 sections for right and left hemispheres from three or more mice for each condition. In inducible *SOX10-iCreERT2* mice, cells were counted through the cortical layers 2-3 and hippocampal DG of 4 coronal sections per mouse. Quantification was performed by investigator blinded to the experimental groups. Images were processed using NIS-Elements Imaging Software, Image J (NIH) and Adobe Photoshop CC.

### Human tissue immunohistochemistry

Human cortical tissue of 68 years old male was obtained postmortem at autopsy with the full consent of each family for the UC Davis Medical Center. Tissue was post-fixed in 4% paraformaldehyde for 24 hours and then transferred into 30% sucrose in PBS until equilibrated. Cortical tissue was sectioned in the coronal plane. 14 μm-thick sections were collected on glass slides. Sections were treated for 30 min with 0.1% of Sudan Black. After quick washes with 70% ethanol followed by PBS, sections were incubated for 1 h at RT with blocking solution containing 10% donkey serum and 0.3% Triton-X in PBS. Sections were incubated with primary antibodies goat anti-Olig2 (1:100, R&D AF2418) and rabbit anti-NeuN (1:100, Millipore ABN78) for two days at 4 °C. Then, sections were washed twice with PBS followed by a single wash with 0.05% Tween-20 in PBS for 10 min, and incubated with secondary antibodies, donkey anti-goat Alexa Fluor-594 and donkey anti-rabbit Alexa Fluor-488 (1:400; Life Technologies) for 2 h at RT. Sections were counterstained with DAPI (1:500, Sigma) for 10 min at RT, washed twice with PBS, once with 0.05% Tween-20 in PBS, and mounted with Mowiol mounting medium (Sigma).

Images were acquired on Nikon Eclipse 90i A1 laser scanning confocal microscope. Negative controls omitting primary antibody were performed to confirm specificity of staining. Images were processed using NIS-Elements Imaging Software and Image J (NIH).

### Super resolution microscopy

Super resolution imaging was performed using a Leica SP8 Falcon confocal microscope with a Lighting detector module. A z-stack of 1320 x1320 images was acquired. Processing of images with the Lighting detector module achieved a sub-diffraction limited resolution image of ~120 nm. Images were processed using ImageJ (NIH). Movie was created from the Z-stack series using ImageJ.

### Fluorescent activated nuclei sorting (FANS)

Mice were euthanized using CO_2_ and cortices and deep gray matter were isolated on ice, and flash frozen in liquid nitrogen for further use. Brain tissue was homogenized in 2 ml of Lysis Buffer (10 mM Tris-HCl, pH 7.4, 10 mM NaCl, 3 mM MgCl_2_, and 0.025% NP-40) in a Dounce homogenizer, and incubated on ice for 15 min. The suspension was filtered through a 30 μm filter to remove debris and pelleted at 500 x *g* for 5 min at 4°C. Pellet was washed with 1 ml of Nuclei Wash Buffer (1% BSA in PBS) twice and passed through a 30 μm filter into a 2 ml tube. After centrifugation at 500 x *g* for 5 min, pellet was resuspended in 450 μl Nuclei Wash Buffer and 810 μl 1.8 M sucrose and carefully overlayed on top of 500 μl 1.8 M sucrose in a 2ml tube. Gradient was centrifuged at 13,000 x *g* for 45 min at 4°C to separate nuclei from myelin debris. Supernatant was discarded until 100 μl bottom fraction containing the nuclei. Nuclei fraction was washed with 1 ml Nuclei Wash Buffer and resuspended in 400ul Nuclei Wash Buffer filtered through a 40-μm FlowMi Cell Strainer. Nuclei were counted using hematocytometer and transferred to staining plates (Fisher Scientific, 12565502). After washing with FANS Buffer (PBS plus 1% BSA, 0.1% sodium azide, 2mM EDTA), nuclei were stained with rabbit anti-NeuN Alexa-647 (1:200, Cell Signaling 62994) and goat anti-Olig2 (1:200, R&D AF2418) on ice for 30-60 min. Nuclei were washed and pelleted by centrifugation at 700 x g for 3 x 5 min at 4°C. Nuclei were stained with Alexa-594 Fluor donkey anti-goat (1:1000) in PBS for 30 min to detect Olig2 primary antibody and washed three times. Nuclei were counterstained with DAPI for 5 min at 4°C. After washing, resuspended nuclei in FANS Buffer were analyzed on Attune NxT Acoustic Focusing Cytometer (Life Technologies). Unstained nuclei, single stain controls and stained AbC Total Antibody Compensation beads (Thermo Fisher, A10497) were used for each experiment to determine compensation parameters and gating. Acquired data were analyzed using FlowJo software (TreeStar). Nuclei populations were gated for single events positive for DAPI, NeuN and Olig2. The data are presented as the percentage of nuclei populations. An aliquot of purified and stained nuclei was sorted on BD Aria II flow sorter for NeuN+Olig2+ population. Sorted nuclei were plated on poly-D-lysin coated glass Lab-Tek II chamber slide (Thermo Fisher Scientific) for confocal analysis.

### Statistics

Statistical analysis was performed using GraphPad Prism 7.5. The number of animals per group used for each study are shown in the figures where each point represents an individual animal. P values were determined using unpaired t-test for two groups comparison or One-way ANOVA with Bonferroni post hoc test to compare three and more groups. In all figures, data are represented as mean ± sem. In all cases, probability values less than 0.05 were considered statistically significant. Images were assembled using Adobe Illustrator CC.

## Extended Data

**Extended Data Fig. 1.**
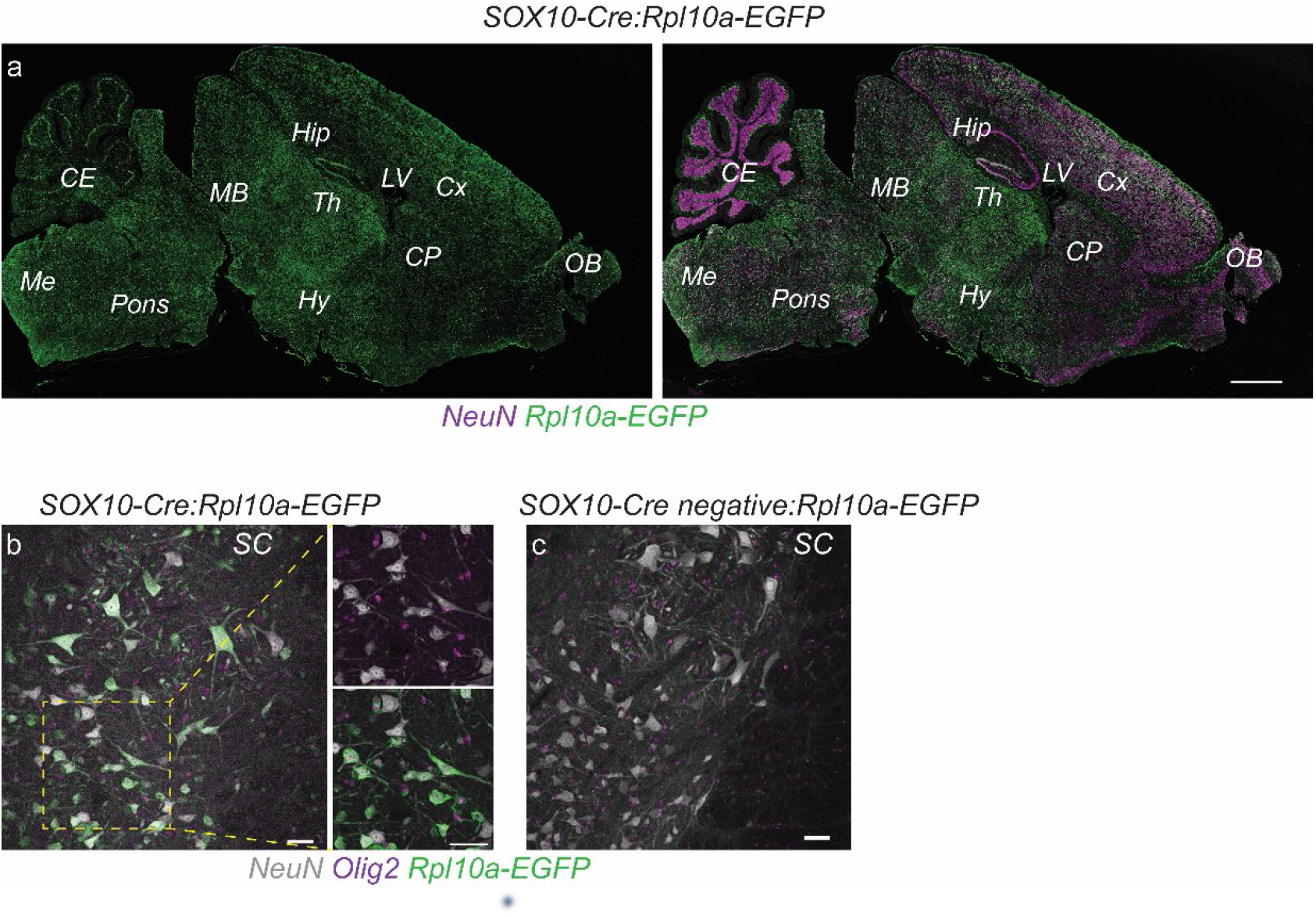
OL-derived material accumulates in neurons throughout the entire CNS. (**a**) Rpl10a-EGFP immunoreactive neurons in the adult brain of *SOX10-Cre:Rpl10a-EGFP* mice. (**b**) Rpl10a-EGFP+ neurons and Rpl10a-EGFP+ OL in the gray matter of the spinal cord in *SOX10-Cre:Rpl10a-EGFP* mice. (**c**) No Rpl10a-EGFP immunoreactivity is detected in the spinal cord of Cre negative siblings of *SOX10-Cre:Rpl10a-EGFP* mice. Scale bars 1000 μm for a; 50 μm for b and c. CE, cerebellum; CP, caudate putamen, Cx, cortex; Hip, hippocampus; Hy, hypothalamus; LV, lateral ventricle; MB, midbrain; Me, medulla; OB, olfactory bulb; SC, spinal cord; Th, thalamus.

**Extended Data Fig. 2.**
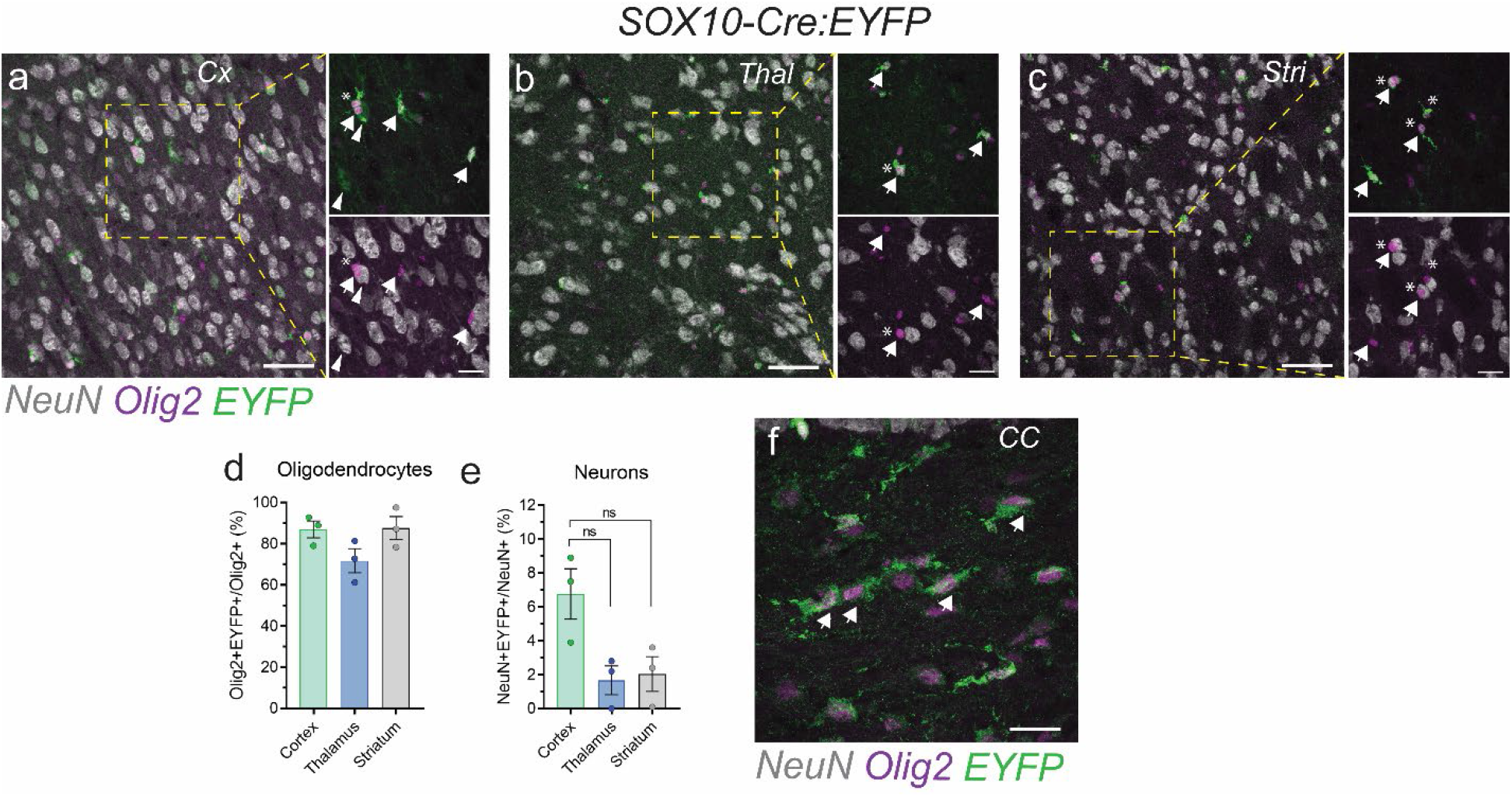
Distribution of unbound EYFP reporter in the adult mouse brain. (**a-c**) EYFP+ OL (arrows) in the cortex, thalamus and striatum of adult *SOX10-Cre:EYFP* mice. Weakly fluorescent neurons in the gray matter of *SOX10-Cre:EYFP* mice (arrowheads). Note satellite OL positioned in close proximity to neuronal soma (asterisks). (**d-e**) Quantification of Olig2+EYFP+ OL and weakly fluorescent NeuN+EYFP+ neurons in the cortex, thalamus and striatum of adult *SOX10-Cre:EYFP* mice. Data are presented as mean ± sem. Each circle represents an individual mouse. *P* values were determined by One-way ANOVA with Bonferroni post hoc test. **(f)** EYFP+ OL in the white matter corpus callosum. Scale bars 50 μm for a-c; 20 μm for a-c enlargements and f. CC, corpus callosum; Cx, cortex; ns, not significant; Stri, striatum; Thal, thalamus.

**Extended Data Fig. 3.**
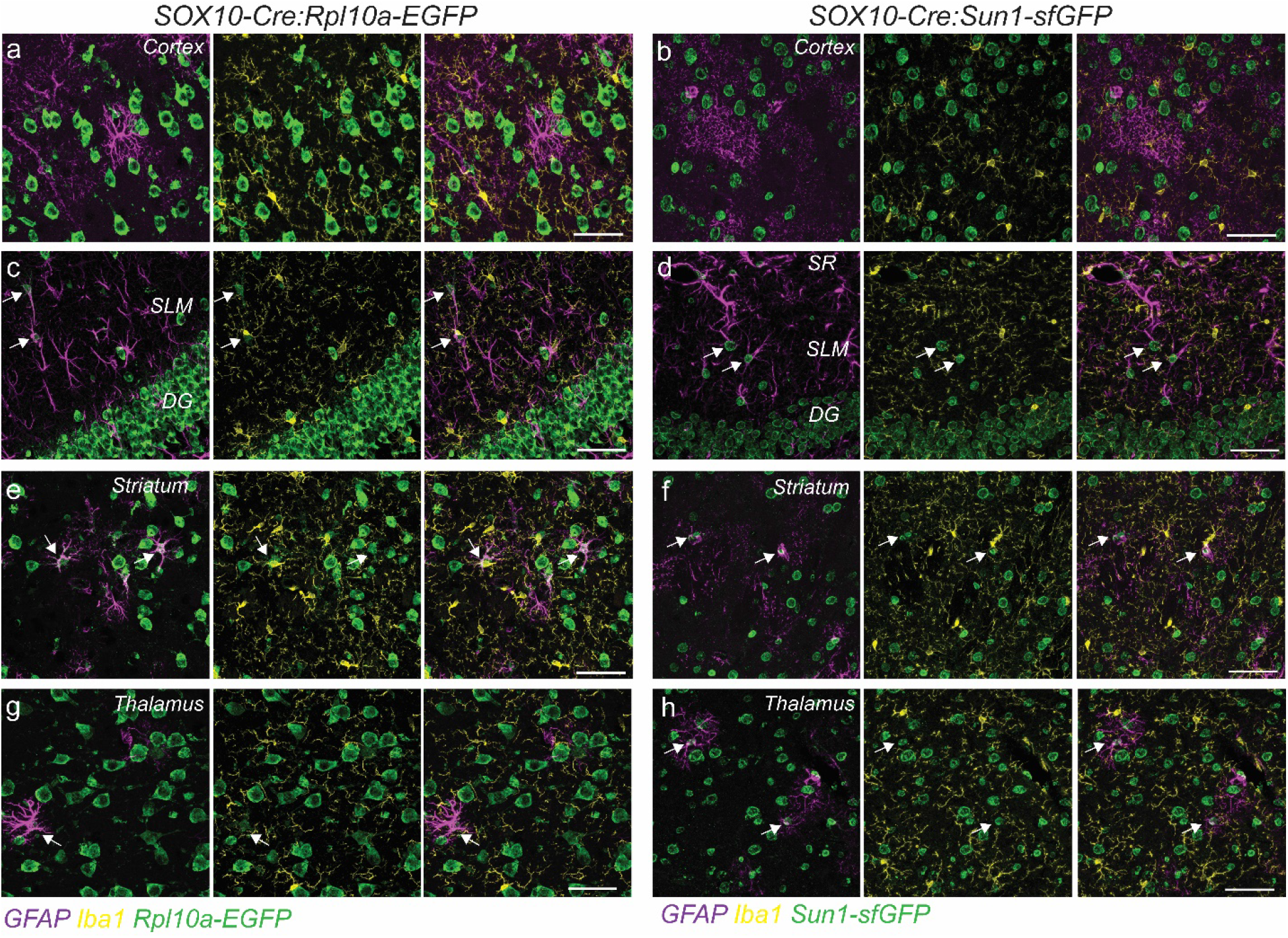
OL-derived ribosomal and nuclear material in astrocytes and microglia in adult mice. (**a-b**) No reporter immunoreactive astrocytes (GFAP+) or microglia (Iba1+) were detected in the cortex of *SOX10-Cre:Rpl10a-EGFP* or *SOX10-Cre:Sun1-sfGFP* mice. (**c-d**) No reporter immunoreactive astrocytes in the DG but some reporter immunoreactive astrocytes were found in the hippocampal stratum lacunosum moleculare (arrows). No microglia containing OL-derived reporters were found in the hippocampus. **(e-h)** Some reporter immunoreactive astrocytes (arrows) were found in the striatal caudate putamen and thalamus. No microglia containing OL-derived reporters were found in the deep gray matter. Scale bar 50 μm for a-h. DG, dentate gyrus; SR, stratum radiatum; SLM, stratum lacunosum moleculare.

**Extended Data Fig. 4.**
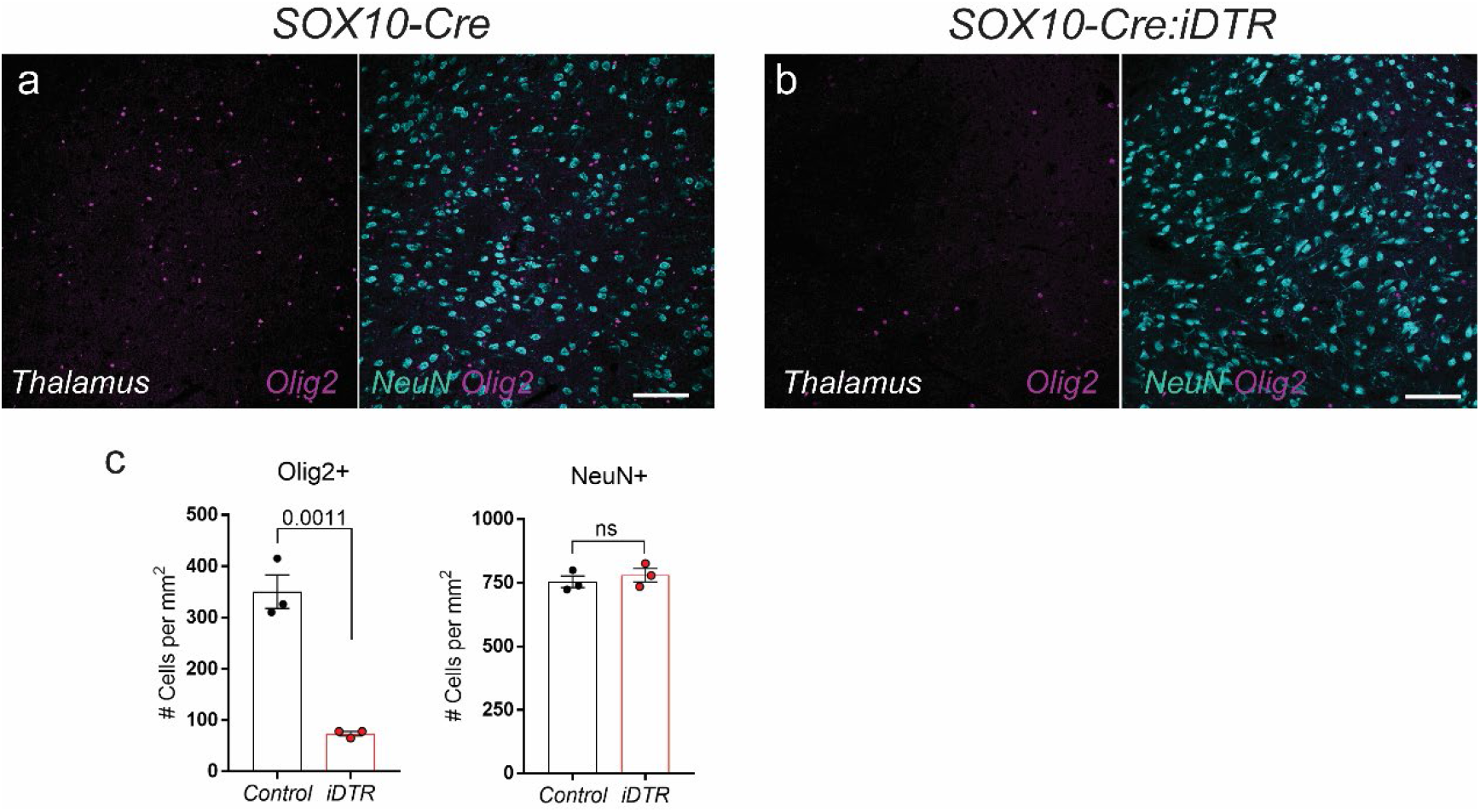
Neurons do not express OL reporter in the thalamus. (**a-b**) *SOX10-Cre:iDTR* and control *SOX10-Cre* mice received stereotaxic injection of 1 ng DT into the right thalamus. Ablation of OL was evident at the injection site in *SOX10-Cre:iDTR* mice 3 days after DT injection. Neuronal density at the injection site in *SOX10-Cre:iDTR* mice was indistinguishable from neuronal density in control *SOX10-Cre* mice. **(c)** Quantification of OL and neurons in the thalamus of *SOX10-Cre:iDTR* and control *SOX10-Cre* mice. Data are presented as mean ± sem. Each circle represents an individual mouse. *P* values were determined by unpaired t-test. Scale bars 100 μm for a-b. DT, diphtheria toxin; DTR, diphtheria toxin receptor; ns, not significant.

**Extended Data Fig. 5.**
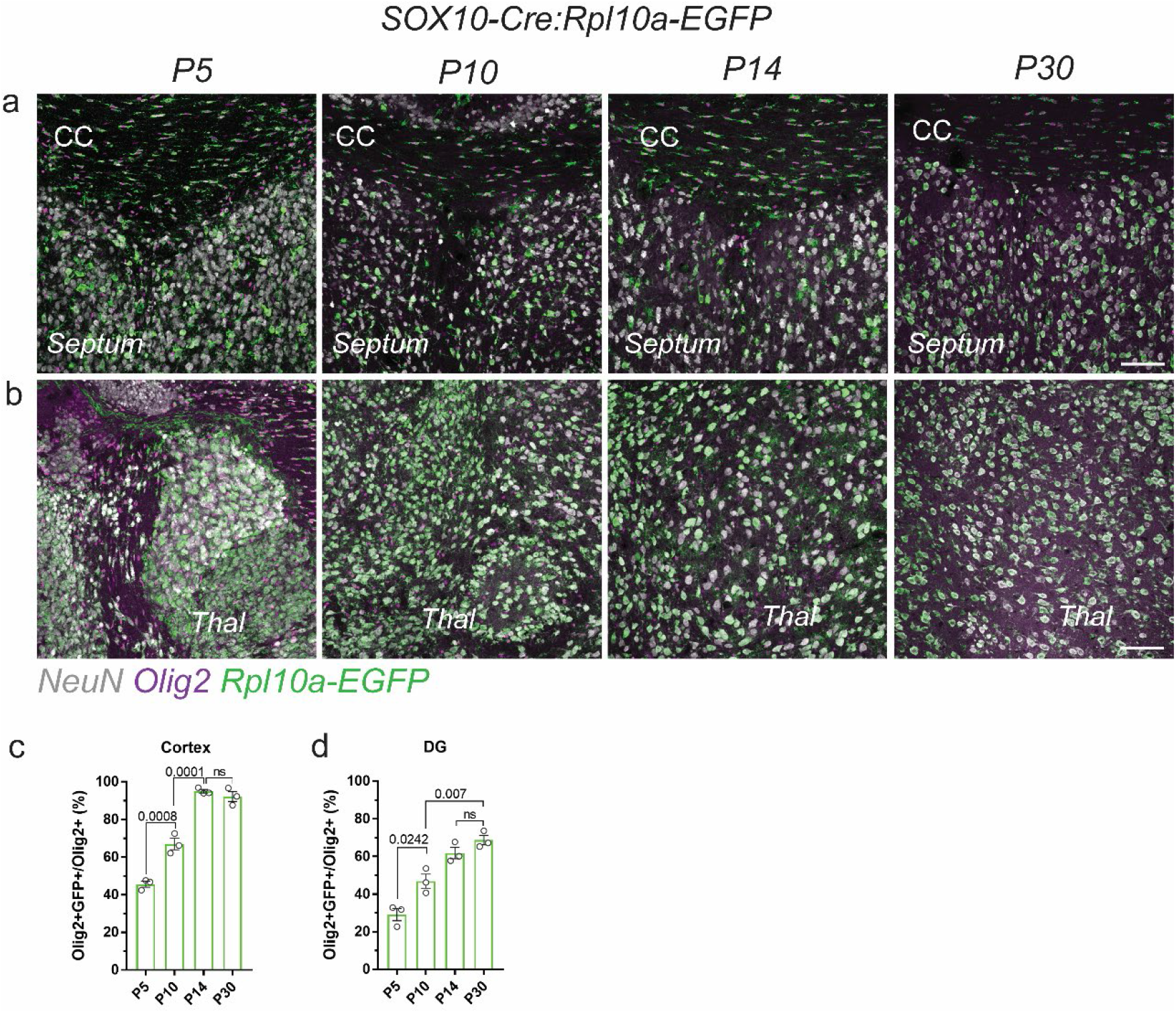
Postnatal propagation of OL-derived material in the mouse brain. (**a-b**) OL-derived reporter protein is found in neurons in the septum and thalamus in *SOX10-Cre:Rpl10a-EGFP* mice early during postnatal development. (**c-d**) Quantification of Olig2+GFP+ OL in the cortex and DG. Data are presented as mean ± sem. Each circle represents an individual mouse. *P* values were determined by One-way ANOVA with Bonferroni post hoc test. Scale bar 100 μm for a-b. CC, corpus callosum; Thal, thalamus.

**Extended Data Fig. 6.**
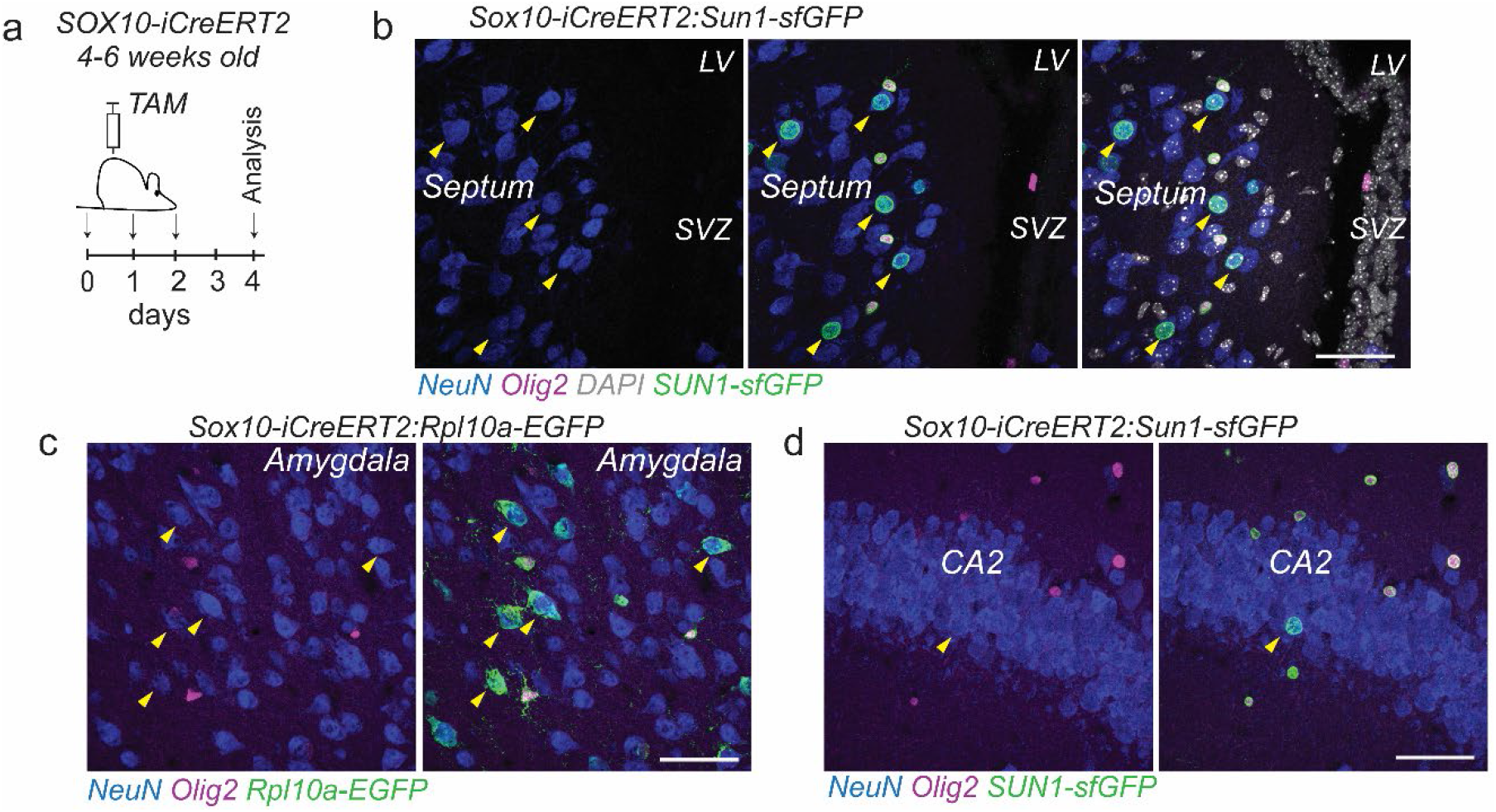
OL-to-neuron material transfer in the septum and amygdala of the adult mouse brain. (**a**) Tamoxifen (TAM) was administered to 4-6 weeks old *SOX10-iCreERT2:Sun1-sfGFP* and *SOX10-iCreERT2:Rpl10a-EGFP* mice. (**b**) At 4 days after first TAM injection fluorescent neurons were found in the septum (arrowheads). (**c**) Rpl10a-EGFP+ neurons (arrowheads) in the amygdala at 4 days post TAM. (**d**) Single neurons immunoreactive for nuclear membrane protein Sun1-sfGFP were found in the hippocampal CA (arrowhead) at 4 days post TAM. Scale bar 50 μm for b-d. CA, cornu ammonis; LV, lateral ventricle; SVZ, subventricular zone.

**Extended Data Fig. 7.**
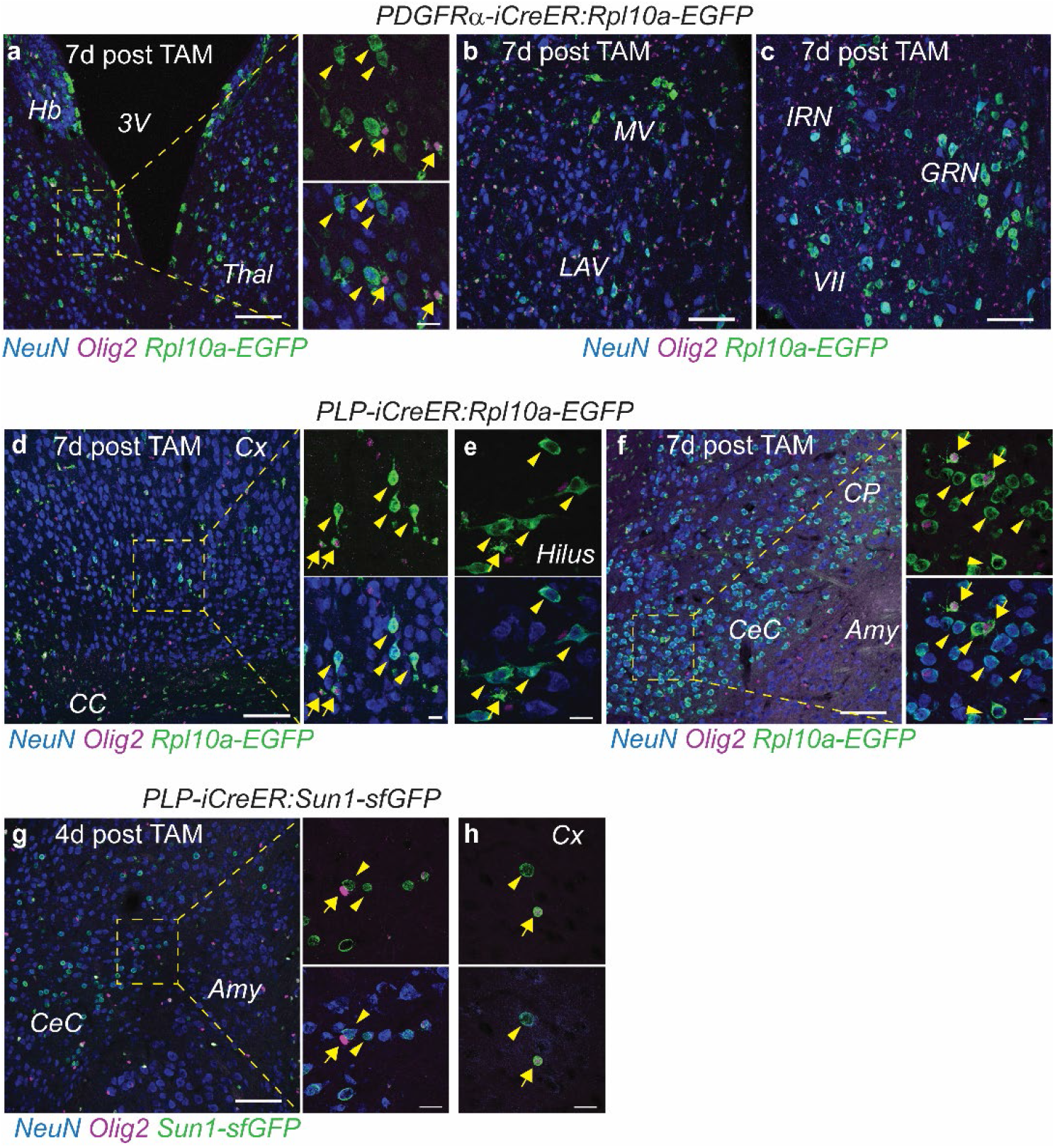
Accumulation of OL-derived nuclear and ribosomal material in *PDGFRα-iCreER* and *PLP-iCreER* mice. TAM was administered to 4-6 weeks old *PDGFRα-iCreER:Rpl10a-EGFP, PLP-iCreER:Rpl10a-EGFP* and *PLP-iCreER:Sun1-sfGFP* mice. (**a-c**) In *PDGFRα-iCreER:Rpl10a-EGFP* mice, in addition to Rpl10a-EGFP+ OL (arrows), multiple reporter-positive neurons (arrowheads) were found in the habenular nuclei, thalamus, and brainstem, including parabranchial, spinal trigeminal, and paragigantocellular nuclei at 7 days post TAM. (**d**-**f**) In *PLP-iCreER:Rpl10a-EGFP* mice, in addition to reporter-positive OL (arrows), Rpl10a-EGFP+ neurons (arrowheads) were found in the cortex, hilus of the DG, and DGM, including septal nuclei, thalamus, striatum, and amygdala. (**g-h**) Sun1-sGFP+ OL (arrows) and reporter-positive neurons (arrowheads) in the lateral capsular division of the central nucleus of the amygdala (CeC) and cortex of *PLP-iCreER:Sun1-sfGFP* mouse brain at 4 days post TAM. Scale bar 100 μm for a-c, d, f, g; 20 μm for enlargements in a, d, f, g and for e, h. Amy, amygdala; CC, corpus callosum; CeC, lateral capsular division of the central nucleus of the amygdala; CP, caudate putamen; Cx, cortex; d, day; DG, dentate gyrus; IRN, intermediate reticular nucleus; GRN, gigantocellular reticular nucleus; LAV, lateral vestibular nucleus; MV, medial vestibular nucleus; OL, oligodendrocytes; TAM, tamoxifen; Thal, thalamus; VII, facial motor nucleus.

**Extended Data Fig. 8.**
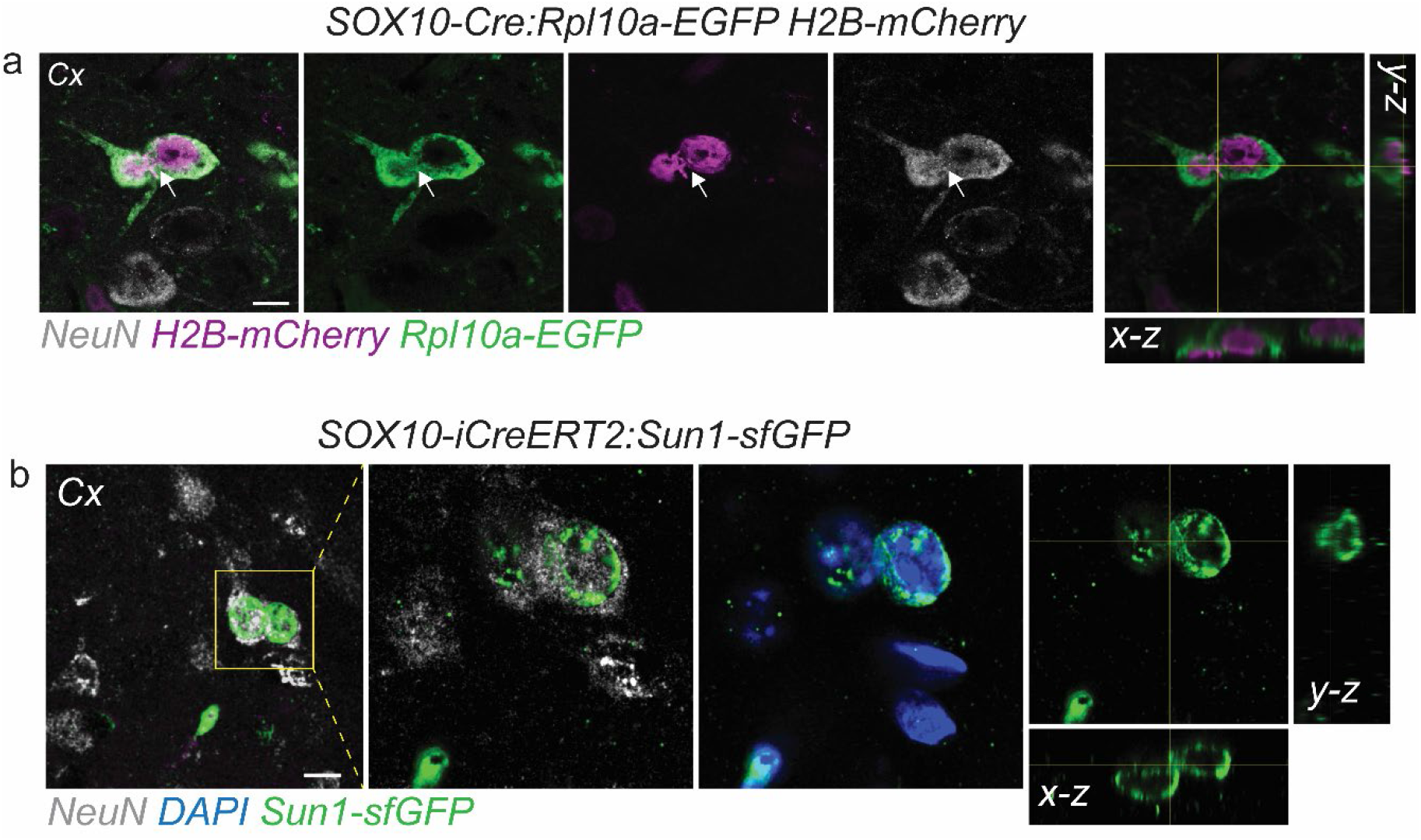
Neuron-neuron internuclear contacts in the adult mouse cortex. (**a**) Two neurons forming internuclear contact (arrow) in the cortex of *SOX10-Cre:Rpl10a-EGFP H2B-mCherry* mouse. Orthogonal projection shows close association of two neuronal nuclei. (**b**) Internuclear contact between two neurons in the cortex of *SOX10-iCreERT2:Sun1-sfGFP* mouse at 4 days post TAM. Orthogonal projection of single plane image shows nuclear membrane contact between two neurons. Scale bar 10 μm for a-b.

**Extended Data Fig. 9.**
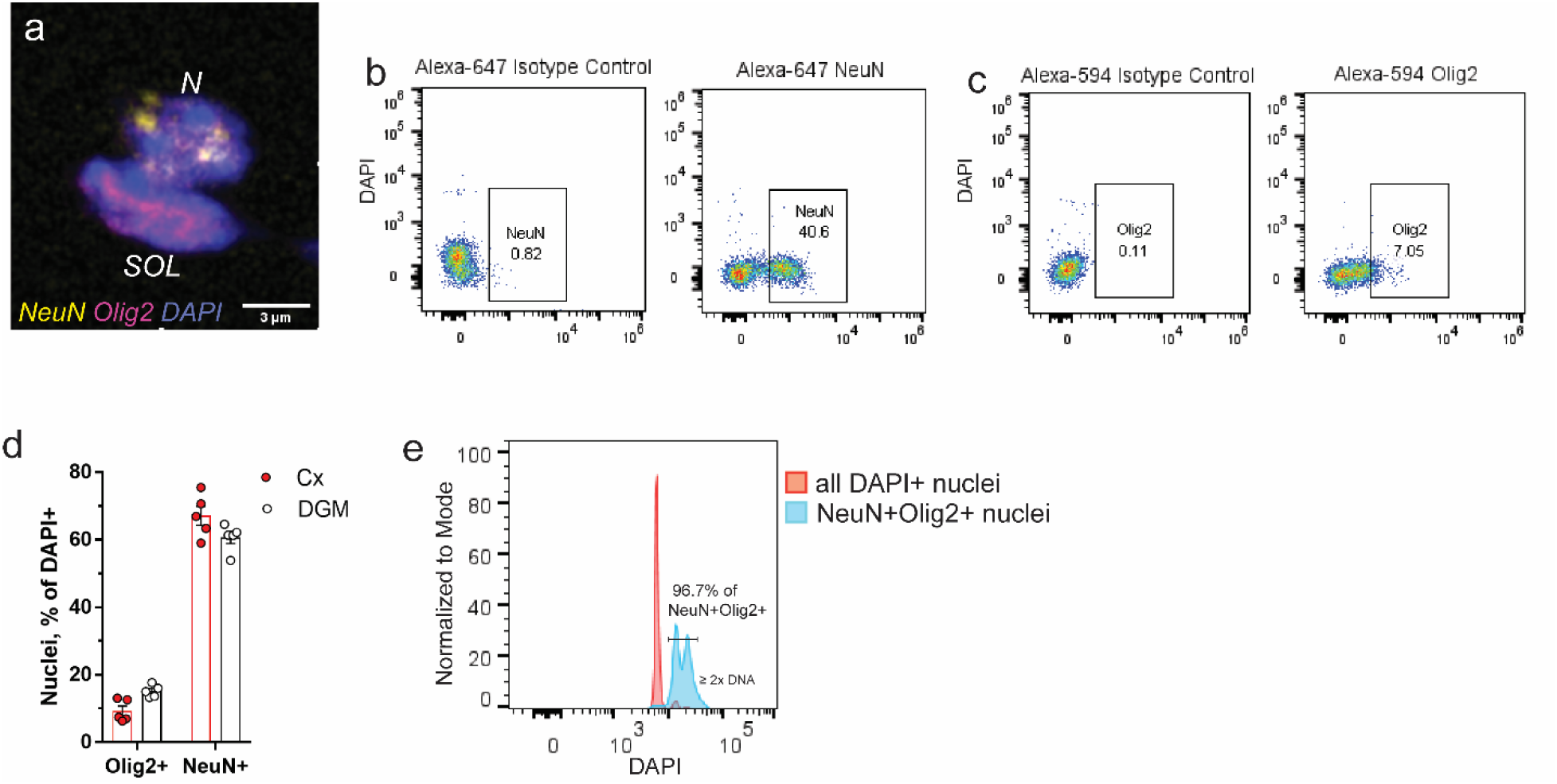
NeuN+Olig2+ double nuclei isolation from the adult mouse brain. Nuclei were isolated from *C57BL/6J* mouse cortex or DGM and sorted by immunofluorescence for NeuN (Alexa-647) and Olig2 (Alexa-594). (**a**) A single plane image shows NeuN+Olig2+ double nuclei after sorting. (**b-c**) Isotype control used for gating NeuN+ and Olig2+ nuclei from DAPI+ population isolated from the mouse brain. (**d**) Quantification of Olig2+ and NeuN+ nuclei isolated from the cortex and DGM. Data are presented as mean ± sem. Each circle represents an individual mouse. (**e**) NeuN+Olig2+ nuclei are enriched in events with 2X and more DNA content. Scale bar 3 μm for a. DGM, deep gray matter; N, neuron; SOL, satellite oligodendrocyte.

